# How increased cognitive load affects the dual-task cost in healthy young adults? A randomized, double-blind sham-controlled study

**DOI:** 10.1101/2021.11.23.469768

**Authors:** Shabnam Behrangrad, Maryam Zoghi, Farshad Mansouri, Shapour Jaberzadeh

## Abstract

Our ability to interact flexibly with the surrounding environment and achieve an adaptive goal-directed response is one of the necessities of balance control. This study aimed to examine the interaction between cognitive demand and the necessity for keeping balance in unstable conditions. We examined the effects of performing two cognitive tasks, namely the Stroop test and Wisconsin Card Sorting Test (WCST), on postural balance in healthy young adults. Stroop and the WCST test assess selective attention and cognitive flexibility in shifting between rules, respectively. Thirty-two healthy adults were included in two experimental conditions (control and treatment) in random order, separated by at least seven days. Standing balance was evaluated by the Sway Medical Mobile application in eyes open (EO) and eyes closed (EC) in different stance positions: feet apart, feet together, semi-tandem, tandem, and single-leg stance (SLS). Balance was evaluated before and after the cognitive test in each experimental condition. Our findings indicate that performing cognitively demanding tasks adversely affected the balance ability in more demanding balance tests such as the SLS with EC (P □ 0.05). However, no significant changes were seen in other balance tests (P □ 0.05). Additionally, no significant changes were seen in balance ability after the Stroop or Wisconsin card sorting test alone. These results confirm that performing cognitively demanding tasks significantly reduced the ability to keep balance in less stable conditions. These findings have significant implications in understanding and preventing falls and incidents resulting from an impaired balance in complex and cognitively demanding conditions.

## Introduction

In real life, there are multiple situations, like facing unexpected obstacles while walking, that we need to withhold or modify our planned behavior before triggering events, such as movement or stimuli from visual, vestibular, or somatosensory systems ^1–3^. In these situations withholding inappropriate or unrequired motor responses and regulating the proper motor responses are necessary to facilitate the execution of chosen goals during motor control ^1–4^. This set of cognitive processes is called executive function ^5–8^. In this study, to simplify for the reader, executive function and cognition are used interchangeably.

Indeed, the executive function enables us to interact flexibly with the surrounding dynamic environment to execute an adaptive goal-directed response ^1–3^. The evidence behind this is found in developmental studies that showed better development of the primary, premotor and supplementary motor areas leads to better implementation of executive functions in the early stages of life and adulthood ^9–19^. Also, studies on children with cognitive problems suggested a direct linear relationship between motor control and cognitive ability ^10–15, 18, 20–30^. Additionally, studies show that role of the executive function is significant in more complex tasks or the tasks needing a higher level of cognitive involvement ^5–8, 31–34^.

The balance tasks can be categorized into single- and dual-tasks, based on the level of cognitive function involved in those tasks. Single-task balance is when the individual performs only one simple balance task at a time ^35, 36^. On the other hand, dual-task balance is an experimental process of performing cognitive and/or motor activities while maintaining a balance task ^35, 36^. The role of executive function is more significant in dual-task activities as the cognitive demand is much higher. Thus, it can be said dual-task paradigm represents more significant mutual communication between motor and executive functions ^31–34^. Previous studies have shown that performing a cognitive interference task such as serial subtraction of numbers, memorizing words, etc., during a balance task reduces the postural control abilities in healthy adults ^10, 33, 37–42^. This decline may result from the overall scarcity of the cognitive resources and, consequently, the allocation of less sufficient cognitive resources during a dual-task balance paradigm ^32^. This interdependence was investigated in more detail in recent years, led to a better understanding of a phenomenon known as cognitive-balance interference ^10, 15, 17, 43^.

Cognitive-balance interference refers to the simultaneous performance of a cognitive task and a balance task, which interferes with the performance of one or both tasks ^44–46^. It evaluates the effects of adding a cognitive task to a single balance task on balance performance. Generally, the result of these studies indicates that cognitive ability and balance are complementary and inseparable, and proper balance maintenance cannot be considered a purely automated sensorimotor activity ^41, 46–50^. The importance of cognitive-motor interference is more evident in daily life activities, mainly when we perform multiple concurrent tasks, such as talking while walking, drinking while driving, cutting something while talking, etc. Performing almost all of the activities of daily living (ADL) requires adequate balance, coordination, critical thinking, and in summary, balance maintenance during the concurrent performance of two or more tasks ^31, 32, 44, 45, 51^.

Cognitive-motor interference can be evaluated by the dual-task cost (DTC) ^52–55^. The DTC is defined as changes in the outcome from single- to dual-task condition (DTC = 100 × [(dual-task score—single-task score)/single-task score]) ^52–55^. The DTC can be positive or negative, and a negative score indicates the lower dual-task ability, whereas a positive score shows higher dual-task ability ^52–55^. Furthermore, sensory information is obtained from the visual, somatosensory, and vestibular systems can affect the postural balance. ^56, 57^. For example, the information acquired by vision is responsible for spatial orientation and body movement relative to the surrounding environment. Therefore, changes in the visual feedback, eyes open (EO) or eyes closed (EC), may provoke different postural control strategies ^56, 57^. Thus, the first aim of this study was to investigate the effects of the cognitive load on the postural balance performance and DTC in different balance tasks with EO and EC.

Additionally, in most ADLs, some conflicts in information processing make us choose between behavioral options to resolve these conflicts. Also, because of the multiple processes encompassed in executive function, several measurements are suggested in the literature to assess it ^58–65^. Ideally, a measure should indicate the initial executive act of inhibiting the dominant response (e.g., reaction time and the tendency to perseverate despite external feedback) and other executive processes called into play. The Stroop test (ST) ^58–61^ and the Wisconsin card sorting test (WCST) ^62–65^ are neuropsychological paradigms to evaluate these conflicts from two different sides of view. The ST measures selective attention ^58–61, 66–68^ and the ability to inhibit an overlearned response in favor of the new one ^66–68^. The WCST assesses the participants’ ability to switch attention from one aspect of an object to another aspect ^69^. This test evaluates abstract reasoning, cognitive strategies shifting ability to achieve a goal and modulating impulsive responding ^62–65^. As discussed, identifying the specific executive domains associated with postural balance changes during dual-task activities may help inform clinical evaluation and treatment planning. However, no study investigated the effects of different executive function tests on postural balance. In addition, research lacks in identifying if postural control is more affected by selective attention (evaluated by ST) or shifting and switching attention (evaluated by WCST) abilities as the two essential components of executive function. Therefore, to answer this need, the second aim of this study was to investigate the effects of the ST ^58–61^ and the WSCT ^62–65^ on single- and dual-task balance with EO or EC in healthy young adults.

## Methods and material

### Participants

In this study, thirty-two healthy volunteers were included from a pool of young, healthy, non-smoking adults aged 18-35 years (16 males, 16 females; mean age 25 years ± 0.14). Participants were recruited using a simple non-probability sampling method. All participants were right-handed, determined by the Edinburgh Handedness Inventory (58.23 ± 8.8) ^70^.

The exclusion criteria were any history of musculoskeletal, neurological, or rheumatoid disorders that might affect balance and cognitive level, and consumption of medications for any neurological condition ^71, 72^. The experimental protocol was done in accordance with the Declaration of Helsinki and approved by the Human Research Ethics Committee, Monash University, Melbourne, Australia. Informed consent was obtained from all participants included in the study.

### Study design

A randomized double-blinded crossover, cross-controlled pre- and post-test study design was utilized in this study. The design includes the participation of volunteers in two experimental conditions in a random order (Figure 1). In study 1, the effects of induced cognitive load by performing two cognitive tasks (ST and WCST) on balance were assessed. In study 2, the effects of induced cognitive load by ST or WCST were investigated separately on postural balance. Each cognitive task was taken about 15 minutes to be performed.

**Figure 1.**
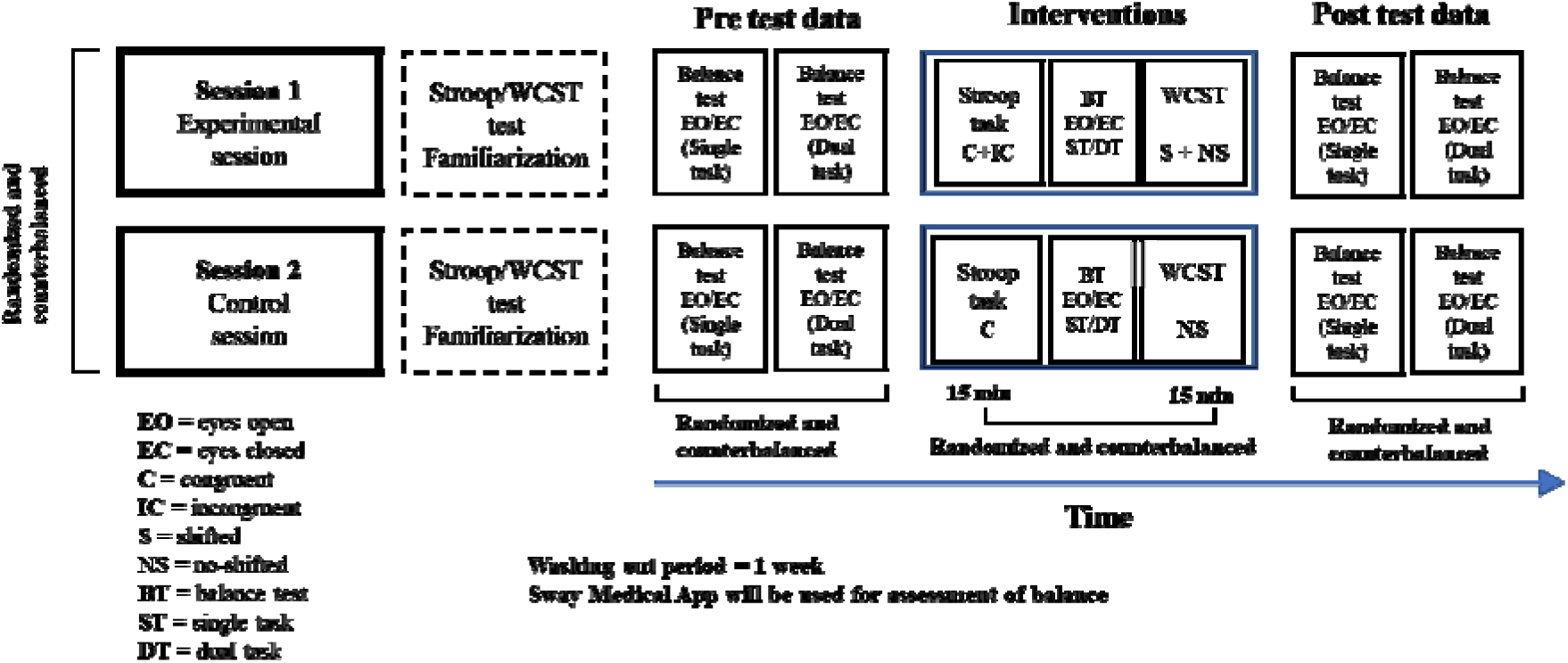
The study set-up. In study 1, the balance was assessed before and after 30 minutes of cognitive tasks (ST and WCST) to investigate the effects of 30 minutes induced cognitive load on balance. In study 2, the balance was assessed before and after 15 minutes of cognitive tests (ST or WCST, not both) to understand the effects of each of these tests on balance.

Also, participants were blinded to the purpose of this study and the experimental conditions and were unaware of the allocated session. Each participant was asked about the nature, experimental or control, of the session they had received at the end of each session to check the blinding integrity. All participants attended two experimental sessions, pseudo-randomly in a counterbalanced manner, separated by at least seven days washing-out period (7.11 days ± 0.65) ^73^. Two researchers were involved in this study, one as the assessor of the outcome measures and the other as the balance and cognition tests administrator. The assessor was in charge of data collection and analysis and was blinded to all experimental conditions and the allocation. The administrator was responsible for balance and cognitive tests performance and not involved in data analysis.

### Experimental procedures

#### Assessment of balance

In this study, the static standing balance was assessed by the Sway Medical Mobile application (LLC, Tulsa, OK, USA). The Sway Medical application is a tool for the assessment of balance. ^74–77^. This application is a phone-based application that generates quantitative scores and helps clinicians and researchers track change over time. It allows the user to quickly and reliably obtain objective, quantifiable balance measurements and evaluate a participants’ initial status and subsequent progress ^74–77^. Studies have shown that the Sway Medical application is a consistent and reliable ^74, 75, 78, 79^, and valid ^74, 76, 80, 81^ tool for evaluating standing balance.

This application is designed to collect rapid, objective evaluation of balance by using built-in mobile sensors. It is also intended to provide healthcare professionals with a quantitative tool for assessing balance ^74–76^. This application is a U.S. Food and Drug Administration-approved mobile device software application installed on a participants’ mobile device. The application accesses all three axes of the device’s accelerometer (anteroposterior, mediolateral, and longitudinal) ^75, 76, 82^. It measures the instantaneous acceleration, which arises in response to postural control and corrective movements. It provides scores ranging from 0 to 100, and higher scores indicate a better balance ^74–76^ (Figure 2).

**Figure 2.**
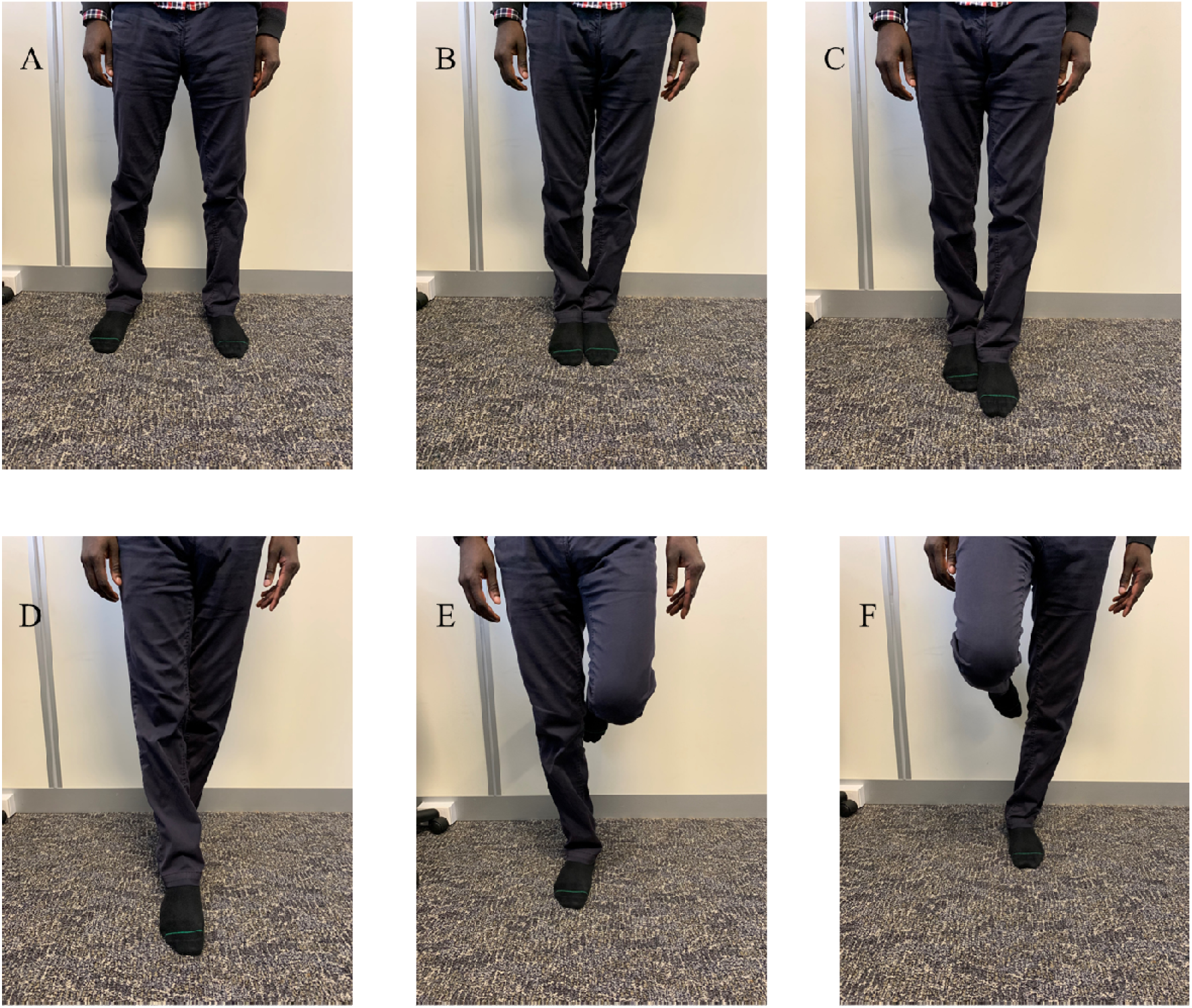
Different stance positions are utilized in the Sway Medical application. A, bipedal (feet apart), B, bipedal (feet together), C, semi-tandem stance (right foot forward), D, tandem stance (right foot forward), E, single-leg stance (right), and F, single-leg stance (left).

In this study, the postural balance was assessed in seven stance positions: 1. bipedal (feet apart), 2. bipedal (feet together), 3. semi-tandem stance (right foot forward), 4. tandem stance (left foot forward), 5. tandem stance (right foot forward), 6. single-leg stance on the right (SLS on Right). 7. single-leg stance on the left (SLS on Left). (Figure 2).

In each stance position, the participant was requested to hold the mobile phone upright over the midpoint of his/her sternum (Figure 2). In all of the balance tests, the first trial was performed as a practice trial. Participants were asked to stand on while holding the phone arms at their sides and were instructed not to move their feet throughout the testing procedure. The balance in these stance positions was assessed with EO and EC for 10 seconds on a firm surface.

Additionally, each test was performed once as a single-task and also as a dual-task. The dual-task was defined as counting backward from a random number between 10 to −10 while the balance test was performed. In contrast, during single-task, the participants have performed the balance test alone. The order of single- and dual-task was counter-balanced in this study, and no instructions were given regarding which task is first.

#### Cognitive tests

Two established tests of executive functions, the Stroop test (ST) ^58–61^ and WCST ^62–65^, have been used in this study. Both tests were carried out using the Psytoolkit online testing platform. Psytoolkit is a free-to-use toolkit for programming and running cognitive tests and surveys ^83–89^. The ST and WCST tests will be explained in the following sections:

#### Stroop test

The ST was used to evaluate the resolving ability of the conflict that happened in information processing ^58–61^ and flexibility to shift cognitive set ^66–68^. It’s been shown when such a conflict occurs between behavioral choices, the speed and accuracy of performance are adversely affected. This effect is referred to as the ‘conflict cost’^58–61^. It has been shown that the computerized ST is a reliable ^66, 90–94^ and valid ^66, 91, 95, 96^ test to measure selective attention capacity and processing speed.

In this test, two types of stimuli were used: 1) congruent stimuli: the name of the color and the actual ink color are the same, and 2) incongruent stimuli: the name of the color and the actual ink color are not the same. The participants were asked to respond to the color they saw, not words they read during this test. Each testing session includes 100 trials (about 15 minutes). The experimental group received a mix of congruent and incongruent trials, whereas, in the control group, participants only received congruent trials. Before performing the actual test, participants received the instruction and conducted a 20-trial (all congruent) to become familiar with the test.

#### Wisconsin card sorting test

The WCST evaluates the participant’s ability to shift attention from one aspect of an object to another ^69^and modulate cognitive strategies toward achieving a goal ^62–65^. In this test, a set of four cards is presented to the subject in each trial, containing one to four equal figures represented by one of four possible shapes, with one of four possible colors (Figure 4). The participant is requested to match the shown card to one of those four cards based on the similarity in color, shape or number of elements without being instructed on the sorting rule. Also, this sorting rule changed without notice during the test, and the participant had to modify the response to feedback based on such changed rules. Correctness of the response is then given according to their choice (“correct” for a correct response, “wrong” for a wrong response and “too slow” if the participant didn’t respond in a specific amount of time).

The WCST included 100 trials with unexpected changing sorting rules in the experimental group; and one changing sorting rule in the control group. Thus, three tests were developed for each matching rule and used randomly for the control group. Before the test, participants received instruction and performed a 20-trial familiarization session (all with one sorting rule) ^62–65^.

### Data analysis

The data were analyzed by SPSS version 22 (IBM Corp., Armonk, NY, United States). A one-way repeated measure ANOVA (RM-ANOVA) was performed on baseline values in different experimental conditions to rule out the carry-over effects of experimental conditions. In addition, the normal distribution of data for each outcome measure was evaluated by the Shapiro-Wilk test. In study 1, a two-way RM ANOVA was carried out to assess the effects of cognitive task performance on DTC. The two independent variables were “cognitive task difficulty” with two levels (experimental and control conditions) and “time” with two levels (T_pre_, T_post_).

In study 2, a three-way RM ANOVA was carried out to evaluate the effects of the cognitive test (ST and WCST) on balance, with two independent variables, “the cognitive task difficulty” with two levels (experimental and control conditions) and “time” with two levels (T_pre_, T_post_) and one dependent variable (performing ST or WCST as a cognitive task).

In all analyses, Mauchly’s test was performed to assess the sphericity assumption validity for RM ANOVA. Also, Greenhouse-Geisser corrected significance values were utilized when sphericity was lacking ^97^. The post-hoc comparisons using the Bonferroni correction were carried out when ANOVA showed significant results. The critical level of significance was set to P □ 0.05. All results are displayed as Means ± Standard Deviation (SD). All of these tests were applied on DTC. As mentioned in the introduction, the DTC is the level changes in the outcome from single- to dual-task condition and is calculated by this formula (DTC = 100 × [(dual-task score—single-task score)/single-task score]) ^52–55^.

## Results

All participants completed both experimental sessions. The Shapiro-Wilk test showed normality in all data sets. The results of the one-way RM-ANOVA showed no significant difference in baseline values in any one of the experimental conditions. (Table 1) (p □ 0.05).

**Table 1.** Baseline balance measurements. Means ± Standard Deviation (SD). FA: feet apart, FT: feet together, US: unilateral stance, Rt: Right, Lt: Left, EO: eyes open, EC: eyes closed. P value< 0.05 is considered significant. Data are reported as mean ± SD. Lines show the means.

| Baseline measurements | Dual-task cost<br>(Control) | Dual-task cost<br>(Condition) | F value | P value |
| --- | --- | --- | --- | --- |
| FA EO | 1.9 ± 16.86 | 1.37 ± 6.5 | 0.27 | 0.6 |
| FT EO | 0.36 ± 5.55 | -0.81 ± 2.74 | 1.13 | 0.29 |
| Semi-tandem EO | 0.3 ± 7.34 | 0.57 ± 3.16 | 0.04 | 0.83 |
| Tandem stance (R forward) EO | -1.41 ± 6.74 | -0.82 ± 7.86 | 0.1 | 0.75 |
| US on Lt EO | -1.86 ± 6.86 | -2.13 ± 6.44 | 0.89 | 0.34 |
| US Rt EO | -1.9 ± 7.79 | -2.3 ± 9.8 | 0.16 | 0.68 |
| FT EC | -0.55 ± 5.33 | 2.82 ± 14.55 | 1.42 | 0.23 |
| Tandem stance (Rt forward) EC | -2.03 ± 12 | -0.24 ± 8.72 | 0.46 | 0.5 |
| Tandem stance (Lt forward) EC | -1.69 ± 11.43 | -0.89 ± 12.14 | 0.07 | 0.79 |
| US Rt EC | -7.89 ± 14.83 | -7.36 ± 19.6 | 0.01 | 0.9 |
| US Lt EC | -2.86 ± 8.68 | -0.34 ± 6.73 | 1.61 | 0.2 |

### Study 1: The effects of increased cognitive load on dual-task cost

A two-way RM ANOVA was performed to evaluate the effects of cognitive task performance on DTC with two independent variables, “cognitive task difficulty” with two levels (experimental and control conditions) and “time” with two levels (T_pre_, T_post_). If the time factor is significant in this analysis, the increased cognitive load significantly affects postural balance. Likewise, if the cognitive task difficulty is significant, the level of difficulty in the cognitive load affects the postural balance significantly. The two-way RM-ANOVA showed no significant effect of task difficulty in any one of the balance testing. There was also no significant effect of time on the DTC in any balance tests, except in the SLS on the Left with EC (Table 2 and 3) (Figure 3). There was also a significant interaction between time and cognitive task difficulty factors when applied on the DTC in the ‘SLS on the left side of the body with EC’. However, there was no significant interaction between these factors for the other balance tests (Table 2 and 3) (Figure 3).

**Table 2.** A summary of the results for the two-way RM-ANOVA for assessment of cognitive load on dual-task cost in eyes open (feet apart, feet together, semi-tandem stance, tandem stance with left foot forward, tandem stance with right foot forward, single-leg stance on the right, and single-leg stance on the left). FA: feet apart, FT: feet together, US: unilateral stance, Rt: Right, Lt: Left. P value< 0.05 is considered significant.

|  | FA |  | FT |  | semi-tandem stance |  | tandem stance |  | US on Rt |  | US on Lt |  |
| --- | --- | --- | --- | --- | --- | --- | --- | --- | --- | --- | --- | --- |
|  | (df, error df)<br>= F value | P value | (df, error<br>df) = F<br>value | P value | (df, error<br>df) = F<br>value | P value | (df, error<br>df) = F<br>value | P value | (df, error<br>df) = F<br>value | P value | (df, error<br>df) = F<br>value | P value |
| Difficulty | (1, 31) =<br>1.182 | 0.286 | (1, 31) =<br>0.977 | 0.331 | (1, 31) =<br>0.016 | 0.899 | (1, 31) =<br>0.713 | 0.405 | (1, 31) =<br>0.1 | 0.754 | (1, 31) =<br>1.306 | 0.262 |
| Time | (1, 31) =<br>2.633 | 0.115 | (1, 31) =<br>0.936 | 0.341 | (1, 31) =<br>0.719 | 0.403 | (1, 31) =<br>0.599 | 0.455 | (1, 31) =<br>0.192 | 0.664 | (1, 31) =<br>0.701 | 0.409 |
| Difficulty<br>and time | (1, 31) =<br>0.806 | 0.377 | (1, 31) =<br>0.08 | 0.779 | (1, 31) =<br>0.028 | 0.868 | (1, 31) =<br>0.78 | 0.384 | (1, 31) =<br>0.088 | 0.768 | (1, 31) =<br>0.717 | 0.404 |

**Table 3.** The results of two-way RM-ANOVA that were performed to assess cognitive load on Dual-task cost in eyes closed have been shown. FT: feet together, US: unilateral stance, Rt: Right, LT: left. P value< 0.05 is considered significant.

|  | FT |  | tandem stance (Rt) |  | tandem stance (Lt) |  | US on Rt |  | US on Lt |  |
| --- | --- | --- | --- | --- | --- | --- | --- | --- | --- | --- |
|  | (df, error df)<br>= F value | P value | (df, error df)<br>= F value | P value | (df, error df)<br>= F value | P value | (df, error df)<br>= F value | P value | (df, error df)<br>= F value | P value |
| Difficulty | (1, 31) =<br>2.664 | 0.113 | (1, 31) =<br>0.045 | 0.825 | (1, 31) =<br>0.002 | 0.964 | (1, 31) =<br>0.756 | 0.392 | (1, 31) =<br>1.586 | 0.218 |
| Time | (1, 31) =<br>0.04 | 0.844 | (1, 31) =<br>0.002 | 0.965 | (1, 31) =<br>0.001 | 0.973 | (1, 31) =<br>1.201 | 0.282 | (1, 31) =<br>4.084 | 0.043 |
| Difficulty<br>and time | (1, 31) =<br>0.371 | 0.547 | (1, 31) =<br>0.303 | 0.586 | (1, 31) =<br>0.073 | 0.789 | (1, 31) =<br>0.179 | 0.675 | (1, 31) =<br>4.706 | 0.039 |

**Figure 3.**
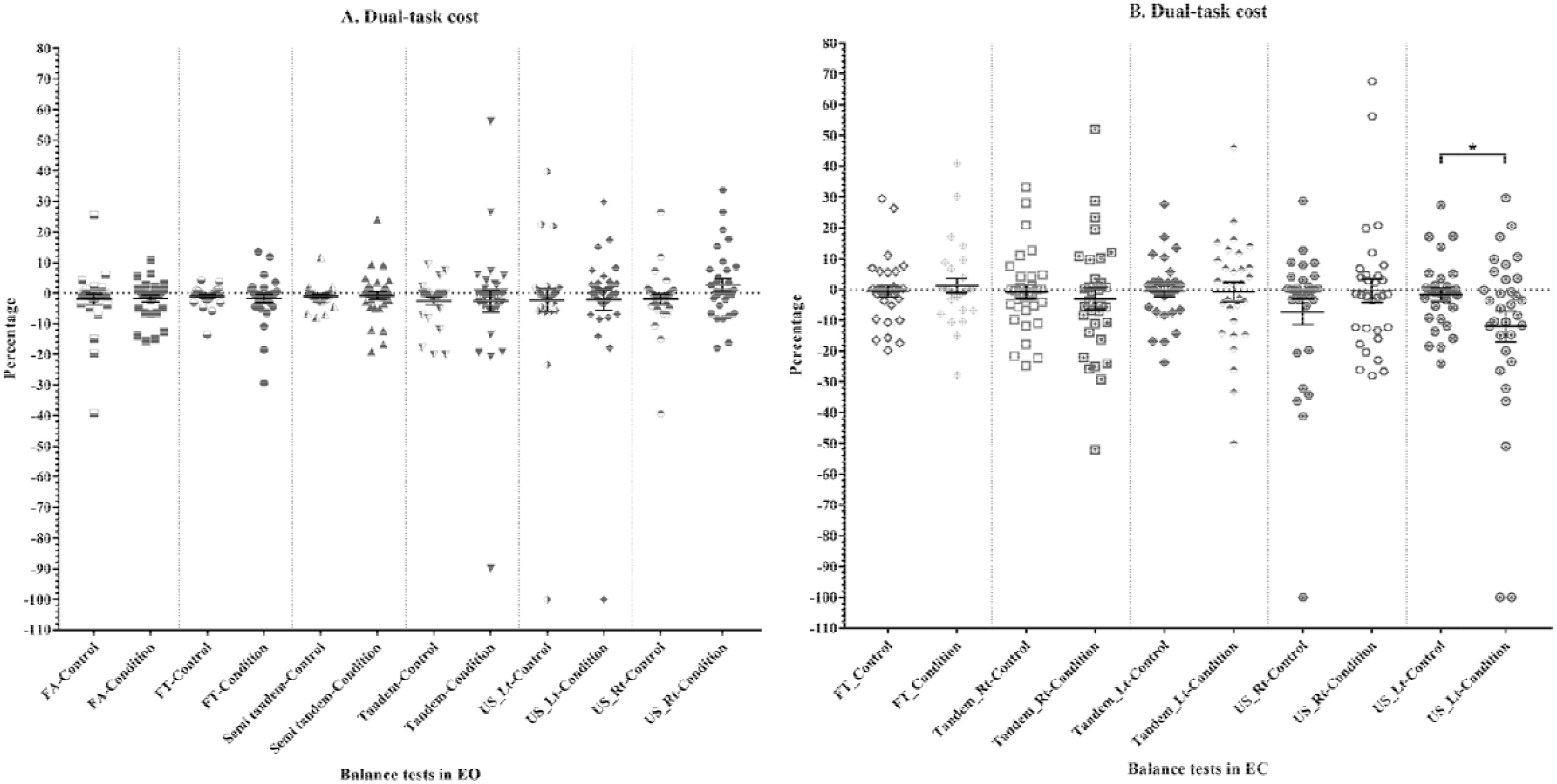
Comparison of the effects of cognitive task performance on different balance tasks in experimental and control groups. A) Dual-task cost in eyes open, B) Dual-task cost in eyes closed. FA: Feet apart, FT: feet together, US: unilateral stance, Rt: Right, LT: left, EO: Eyes open, EC: Eyes closed. P-value < 0.05 is considered significant. Each dot represents the mean value for each participant. Data are reported as mean ± SD. Lines show the means. Error bars indicate SD.

**Figure 4.**
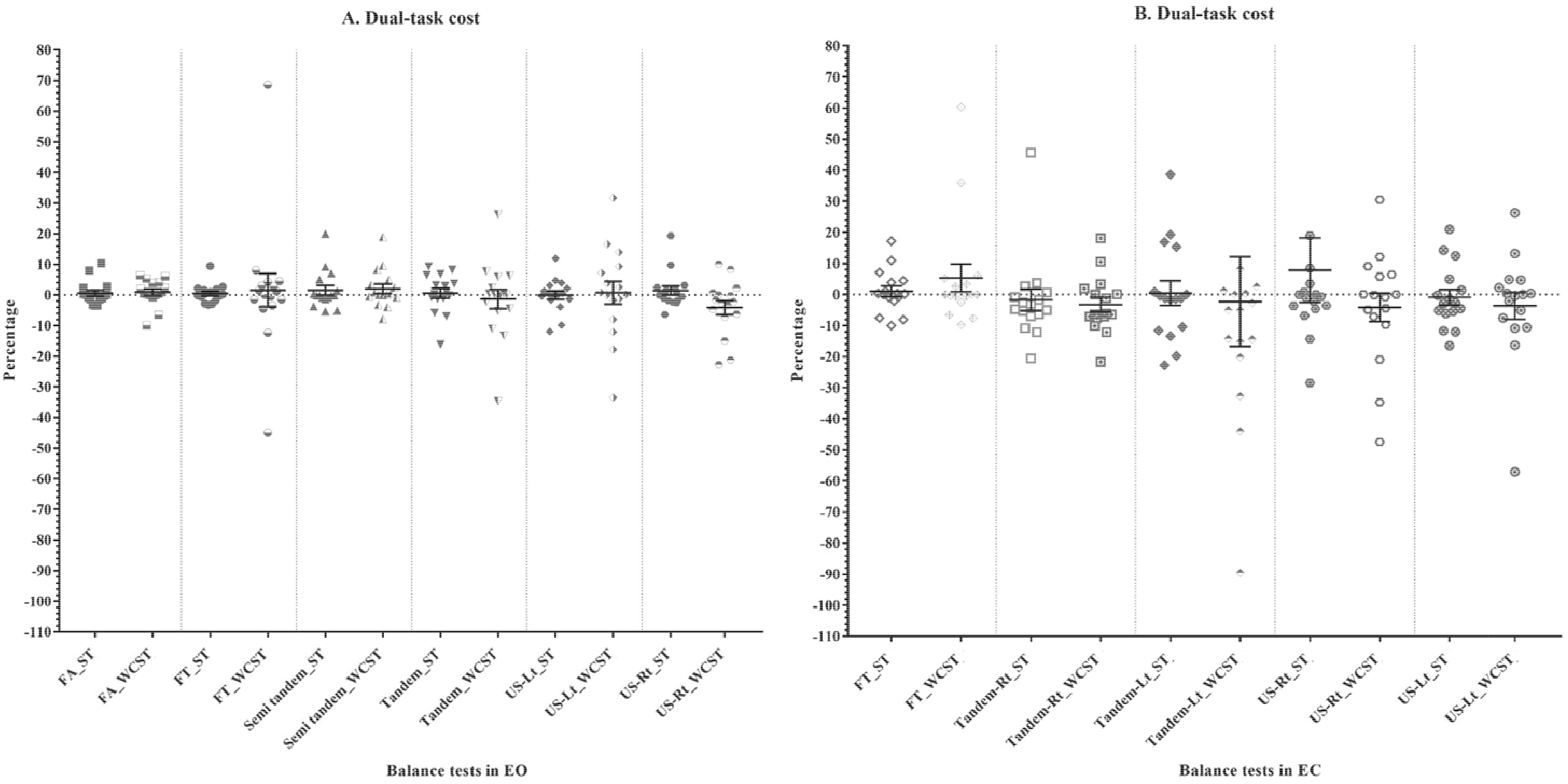
Comparing the effects of the Wisconsin card sorting test and the Stroop test on balance scores in different balance conditions for the experimental and control groups. A) Dual-task cost in eyes open, B) Dual-task cost in eyes closed. FA: Feet apart, FT: feet together, US: unilateral stance, Rt: Right, LT: left, EO: Eyes open, EC: Eyes closed. Each dot represents one participant. Data are reported as mean ± SD. Lines show the means. Error bars indicate SD.

Notably, a significant difference was found in DTC between the control and experimental group in the SLS on Left with EC (p = 0. 04) (Effect size = 0.48, Cohen’s d = −0.97, 0.02), suggesting the performing two cognitive tasks may affect the cognitive-motor interference ability in more complex balance tasks like the SLS on Left with EC in healthy adults (Figure 3).

### Study 2: The effects of induced cognitive load using Stroop test and Wisconsin card sorting test on dual-task cost

A three-way ANOVA was performed to examine the effect of cognitive task difficulty, time and type of executive function test on the balance performance. Two independent variables, “the cognitive task difficulty” with two levels (experimental and control conditions) and “time” with two levels (T_pre_, T_post_) and one dependent variable (performing ST or WCST as a cognitive task) were used in this analysis. The significance in time means the postural balance is affected after an increased cognitive load increased by WCST or ST, and the significance in the cognitive task difficulty suggesting postural balance changes between the two levels of cognitive task difficulty utilized in this study. There was no significant three-way interaction in all of the experimental conditions and time [F _FA, EO_ (1, 31) = 0.784, p _FA, EO_ = 0.384, F _FT, EO_ (1, 31) = 0.754, p _FT, EO_ = 0.392, F _Semitandem, EO_ (1, 31) = 0.739, p _Semitandem, EO_ = 0.397, F _Tandem, EO_ (1, 31) =0.231, p _Tandem, EO_ = 0.634, F _SLS on Left, EO_ (1, 31) = 0.431, p _SLS on Left, EO_ = 0.516, F _SLS on Right, EO_ (1, 31) = 0.988, p _SLS on Right, EO_ = 0.328, F _FT, EC_ (1, 31) = 2.191, p _FT, EC_ = 0.15, F _Tandem on Right, EC_ (1, 31) = 533, p _tandem on Right, EC_ = 0.471, F _Tandem on Left, EC_ (1, 31) = 2.812, p _Tandem on Left, EC_ = 0.105, F _SLS on Right, EC_ (1, 31) = 0.599, p _SLS on Right, EC_ = 0.445, F _SLS on Left, EC_ (1, 31) = 0.051, p _SLS on Left, EC_ = 0.823]. These findings indicate that performing the individual tasks (Stroop or WCST) did not affect the postural control in healthy adults.

## Discussion

### Study 1: The effects of increased cognitive demand on balance

Our findings indicate that performing two executive control tasks affected participants’ balance only in the SLS on the Left with EC condition. This observation suggests that cognitive demand (performing both ST and WCST) only affects balance in more unstable conditions (such as SLS on Left with EC). This finding can be described by the capacity-sharing theory ^98, 99^. According to this theory, to affect the postural balance, the elicited attentional demands during the task should be exceeded central processing capacity ^100–102^, which seems to happen in the SLS on the Left with EC and not the other balance tests. Therefore, that model can explain the balance deterioration found in more complex balance tests like the DTC of the SLS with EC. Also, it is recommended that in healthy adults, the lower dual-task ability can be compensated by different mechanisms in balance like higher toe clearance in gait ^103^ and decrease the speed of movement ^104^, which may be another reason for not having significant balance differences found in other balance tasks in this study. In other words, it seems that the other balance tasks were not challenging enough for the participants’ postural control ability to elicit the hypothesized interaction effects. Therefore, it is suggested that the extra challenge should be added to these balance tasks and/or more complex cognitive tests.

The findings in the current study are in line with other studies investigating the relationship between balance and increased cognitive load ^103–106^. For example, Worden et al. performed a dual-task test with more central interference by adding a more complex cognitive task (visual ST) and obstacle during the balance test. It’s been found the lower dual-task abilities happened when the participants had to switch gaze between the obstacle and Visual ST ^104^. Also, it’s been shown in healthy adults; the lower dual-task abilities happen in more challenging balance tasks like the absence of accurate and consistent visual information in walking ^103^. Likewise, Anderson et al. found that performing cognitive tasks led to lower balance performance and lower cognitive scores obtained when the balance was challenged by external perturbation ^105^. Moreover, Zhang et al. found additional cognitive demand can compromise unpredicted motor control perturbation during walking ^106^.

Furthermore, it’s been suggested that there is a significant interaction between vision and cognitive load on the center of pressure changes in balance tasks ^102, 106–108^. As a result, more attentional resources are involved in processing the executive function load when visual information is occluded in balance tasks, like the SLS on Left with EC. Thus, such situations lead to reducing postural balance control automaticity and decreased balance performance consequently.

The findings in the current study can be explained by the role of the prefrontal cortex in dual-tasking ^109, 110^. It’s been suggested that increased prefrontal cortex activity resulting from dual-task activities may lead to reduced available resources for visual and vestibular processing, causing alterations in shared attentional resources ^109, 110^. Also, it’s been shown that performing prolonged mentally-demanding tasks causes changes in the level of prefrontal cortex electroencephalography activity and increased beta-frequency band power that seems to be related to decreased brain alertness and arousal levels ^111, 112^. Additionally, although maintaining a stable upright posture requires minimal attention, it’s been found that the executive function load can decrease concentration on and attention to the given task ^112, 113^. As well, it’s been suggested higher executive function load may temporarily impair cognitive functioning and, consequently, may increase gait variability and postural sway in healthy adults ^112, 113^.

So far, not many studies have explored the effect of balance conditions on cognitive and motor performance ^114, 115^. Among the few studies in this area, Pellecchia et al. looked at the effect of dual-task training on postural sway in quiet standing on a compliant surface in healthy young adults and suggested dual-task practice improves dual-task performance ^114^. In contrast to the findings in the current study, significantly better motor task performance was seen in the dual-task training group. However, postural sway significantly increased in no-training and single-task training groups.

Similarly, Beurskens et al. evaluated cognitive and motor task performance under single- and dual-task conditions ^115^. Unlike our study, Beurskens et al. found significant longer time in balance and improved standing balance performance in standing. One of the main differences was in the balance tests was used. In Pellecchia et al. and Beurskens et al., only a quiet standing static balance task was used; instead, a variety of more complex static balance tests were utilized in our study to change the proprioception level and visual information participants receive. As it’s been shown visual information is an essential element of postural balance, it is believed that different neuromuscular mechanisms are responsible for regulating static balance control with and without visual input. Also, the other major difference was the training process performed in these two studies, which might be the main reason for increasing balance task automatization compared to the present study and consequently balance improvement in dual-task.

### Study 2: The effects of induced cognitive load using ST and WCST on dual-task cost

This study didn’t find any significant difference between the effects of ST and WCST on DTC. The main reason can be the included participants in this study. As expected, young, healthy adults have proper attentional and balance abilities. Thus, more complex or prolonged tasks are needed to lead to cognitive fatigue to determine how these tests affect balance. Also, the other reason is probably due to healthy intact neural systems; these participants could prioritize their attentional resources to balance and thus could keep their proper balance after each of these tests ^110^. Also, it can be explained by the psychophysiological response to cognitive workload during dual-task ^116^. In the following section, some studies that found different results are evaluated more specifically for each test.

#### The effects of induced executive function load using Stroop test on balance

Few studies evaluated the effects of ST on postural balance and found different results than our study ^115, 117^. Talarico et al. assessed the effects of ST as a selective attention test on squat movement and postural control performance. Interestingly, this study found that the ST significantly leads to a significant sway speed increase in dual-task compared with single-task as the speed of a squat increases ^117^. One of the main differences that may lead to different results can be the balance test selection. The squat test is a functional task with a dynamic center of pressure changes. Thus it can utilize participants’ attentional resources much more than the balance tests selected in the current study. It is believed that cognitive load is added while performing a functional task like squat at a faster speed, increasing the demands on the postural control system to maintain stability. This redirects attentional demands to assist in completing the cognitive task; thus, less effort is directed at the functional task; consequently, worse performance and faster sway speed are seen.

Interestingly, Beurskens et al. investigated the effect of consecutive, concurrent auditory ST practice on single- and dual-task dynamic balance task performance in healthy young adults. This study showed performing the ST consecutively before single- and dual-task dynamic standing practice can improve motor task performance ^115^. These findings can be the results of practicing the dynamic balance in young adults and can be because of the facilitatory effects of the auditory ST on learning balance tasks in healthy adults ^98^. The main difference that led to improvement in Bayot et al. study is the practice that was added to the balance test performance, which can be described by the findings of Plummer et al. According to Plummer et al., variability in patterns of cognitive-motor interference can be explained by the task specificity, individual characteristics, and differences in the measured parameters ^118^. In this classification system, the motor facilitation effects can happen based on the task novelty (i.e., the individual’s previous experience with the performance of the task). Also, the other main difference was using the auditory version of the ST rather than the visual version utilized in the current study, which may differ the effects on the postural balance system. It’s been suggested that the visual interference effect of a selective switch tasks (like ST or flanker task) can increase initial motor program errors and prolong step execution time even in young adults ^98, 119, 120^.

#### The effects of increased cognitive load (cognitive set-shifting) on balance

No study was found on the effects of induced cognitive load by the WCST on postural balance. However, one study evaluated the association of mental flexibility, evaluated by WCST, with poor balance and falls in older adults ^121^. Significant associations between cognitive functioning, balance and falls in older people were seen. Furthermore, Pieruccini-Faria et al. found that the group categorized as high-risk fallers with reduced balance control performed worse in the WCST. This was in line with other studies suggesting poor mental flexibility affects the balance ability and increases the risk of falls in older adults ^121–123^. However, Pieruccini-Faria et al. looked at the association between the postural balance performance and WCST results and didn’t investigate the effects of induced cognitive load on postural balance ^121^. The main reason behind the discrepancy in findings lies in the age of participants. It has been shown that the prioritization between the concurrent cognitive task and balance task in dual-task is different in young and older adults ^34, 116, 124, 125^. While young adults could perform both tasks concurrently, older adults prioritize balance rather than cognitive tasks ^124^. Thus, the response to cognitive load is different.

Also, few studies evaluated the effects of cognitive load with similar executive function tests on balance performance ^126, 127^. These studies utilized 90-minute cognitive tests (the AX-continuous performance task), a visual cognitive task that requires sustained attention, response inhibition and error monitoring and evaluated its effects on the postural balance. Both studies found that this amount of cognitive load leads to mental fatigue and impaired balance control, especially in the most challenging postural contexts like standing with EC. These findings suggested the level of cognitive load is an important factor to modulate balance task in young, healthy adults, and performance of WCST in this study took less than 90 minutes (between 5-10 minutes), the amount of induced cognitive load wasn’t enough to affect postural balance in healthy young adults.

### Limitations

The findings in this study should be interpreted in light of several limitations. First, because the data were obtained from healthy young adults, the results may not be extrapolated to other age groups like the older population. Second, gender has not been assessed as a variable, although conducted on female and male participants. Therefore, the results of this study cannot be gender-specific. Third, in this study, only static balance was evaluated, and no dynamic balance assessment was carried out; thus, the findings from this study could not be extrapolated to dynamic balance.

### Suggestions for future research

Future research should address more dynamic balance tasks or balance tasks in more complex environments. It is also suggested to evaluate the effects of the induced cognitive load on older adults. The other suggestion is to use different executive function tests to assess the other essential aspects of the executive function, such as working memory, cognitive control and determining their impact on the balance. Also, as the effects of cognitive load may be different in both genders, new studies are needed to evaluate the gender effects of induced cognitive load on balance.

## Conclusion

These results confirm that the cognitive load can significantly decrease balance ability, especially in more challenging balance tasks, like in the SLS with EC in healthy young adults. Moreover, this study indicates that the induced cognitive load by ST or WCST alone had no significant effects on balance, indicating the amount of cognitive load induced by each of these tests was not enough to affect the postural balance in healthy young adults.

## Author contributions

Conceived and designed study: SB, FM, SJ; Performed data collection: SB; Conducted the analysis: SB; Interpreted the findings: SB; Wrote the Manuscript: SB; Writing and editing of drafts: SB, MZ, FM, SJ.

